# Expression Ratios of the Anti-Apoptotic BCL2 Family Members Dictate the Selective Addiction of KSHV-Transformed Primary Effusion Lymphoma Cell Lines to MCL1

**DOI:** 10.1101/2022.09.07.507059

**Authors:** Daniel Dunham, Prasanth Viswanathan, Jackson Gill, Mark Manzano

## Abstract

Kaposi’s sarcoma-associated herpesvirus (KSHV) causes several malignancies in people living with HIV, including primary effusion lymphoma (PEL). PEL cell lines exhibit oncogene addictions to both viral and cellular genes. Using CRISPR screens, we previously identified cellular oncogene addictions in PEL cell lines, including MCL1. MCL1 is a member of the BCL2 family, which functions to prevent intrinsic apoptosis and has been implicated in several cancers. Despite the overlapping functions of the BCL2 family members, PEL cells are only dependent on MCL1 suggesting that MCL1 may have non-redundant functions. To investigate why PEL cells exhibit selective addiction to MCL1, we inactivated the intrinsic apoptosis pathway by engineering BAX/BAK1 double knockout cells. In this context, PEL cells become resistant to MCL1 knockdown or MCL1 inactivation by the MCL1 inhibitor S63845, indicating that the main function of MCL1 in PEL cells is to prevent BAX/BAK1-mediated apoptosis. The selective requirement to MCL1 is due to MCL1 being expressed in excess over the BCL2 family. Ectopic expression of several BCL2 family proteins, as well as the KSHV BCL2 homolog, significantly decreased basal caspase 3/7 activity and buffered against staurosporine-induced apoptosis. Finally, over-expressed BCL2 family members can functionally substitute for MCL1, when it is inhibited by S63845. Together our data indicate that the expression levels of the BCL2 family likely explain why PEL tumor cells are highly addicted to MCL1. Importantly, our results suggest that caution should be taken when considering MCL1i as a monotherapy regimen for PEL, because resistance can easily develop.

**IMPORTANCE:** Primary effusion lymphoma (PEL) is caused by Kaposi’s sarcoma-associated herpesvirus. We previously showed that PEL cell lines require the anti-apoptotic protein MCL1 for survival, but not the other BCL2 family proteins. This selective dependence to MCL1 is unexpected as the BCL2 family functions similarly in preventing intrinsic apoptosis. Recently, new roles for MCL1 not shared with the BCL2 family have emerged. Here, we show that non-canonical functions of MCL1 are unlikely essential. Instead, MCL1 mainly functions to prevent apoptosis. The specific requirement to MCL1 is due to MCL1 being expressed in excess over the BCL2 family.

Consistent with this model, shifting these expression ratios changes the requirement away from MCL1 and towards the dominant BCL2 family gene. Together, our results indicate that although MCL1 is an attractive chemotherapeutic target to treat PEL, careful consideration must be taken as resistance to MCL1-specific inhibitors easily develops through BCL2 family overexpression.

## INTRODUCTION

Kaposi’s sarcoma-associated herpesvirus (KSHV) establishes lifelong latency in B cells. While KSHV infections are often asymptomatic, they can develop into lymphoproliferative disorders such as multicentric Castleman disease and primary effusion lymphoma (PEL) in people living with HIV (1). PEL is a monoclonal B cell malignancy often found as effusions in body cavities (2). Patients treated with modified etoposide, vincristine, and doxorubicin with cyclophosphamide and prednisone (EPOCH) have a median survival time of 22 months and a three-year survival rate of 47% (3). To date, no cancer-specific treatment is available for PEL.

PEL cell lines exhibit gene expression profiles closest to plasma cells (4–6) with elevated levels of key plasma cell-associated transcripts such as the master transcription factor interferon regulator factor 4 (IRF4) and genes involved in the unfolded protein response and endoplasmic reticulum stress (ER) pathways (4, 7). The majority of PEL tumors (~80%) are also co-infected with the related gammaherpesvirus Epstein-Barr virus (EBV). The presence of EBV appears to improve prognosis in patients (3). Genetic analyses of PEL tumors do not show mutations or translocations found in other hematological malignancies such as *TP53*, *MYC*, *BCL2*, or *BCL6* (2, 8–10) although alterations in fragile site tumor suppressors have been reported (11). Instead, the defining feature of PEL is that all tumor cells are latently infected with KSHV. The latent virus constitutively expresses only a narrow subset of latency genes and at least 20 mature viral microRNAs.

It is thought that constitutive expression of the KSHV latency genes drives the transformation of PEL. Knockdown of the viral cyclin, viral FLICE inhibitory protein, viral interferon regulatory factors 1-3 (vIRF1 to 3), or viral interleukin 6 all led to a decrease in survival of PEL cells in culture (12–16). These latency genes regulate cellular networks to constitutively activate pathways necessary to support the survival and proliferation of these infected cells. This creates “oncogenic dependencies or addictions” to cellular genes, because tumor cells rely on the continuous signaling of these pathways. For instance, addictions to several oncogenes have been reported in PEL cell lines including the constitutive NF-κB signaling, phosphatidylinositol-3-kinase-mTOR signaling, gp130-STAT3 signaling, high expression of the p53 inhibitor MDM2 and the IRF4 transcription factor, and c-Myc stabilization (9, 17–25).

We recently performed genome-wide CRISPR knockout screens to comprehensively identify human oncogenic addictions or cellular genes that are required for the survival and/or proliferation of PEL cell lines (26). Our study confirmed the addictions to well-described pathways above. Importantly, we uncovered new dependencies that are consistently required in eight different PEL cell lines which included genes that are commonly essential regardless of cell type and genes that are strongly selective. Of note, we identify a strong addiction to the myeloid cell leukemia-1 gene (MCL1). MCL1 belongs to the BCL2 gene family, of which there are six members: MCL1, BCL2, BCL-xL (encoded by *BCL2L1*), BCL-W (encoded by *BCL2L2*), A1 (encoded by *BCL2A1*), or Diva (encoded by *BCL2L10*). The BCL2 family functions to inhibit intrinsic apoptosis. In healthy cells, monomers of the apoptotic effector BAX or BAK1 are sequestered on the mitochondrial outer membrane (MOM) by the BCL2 family proteins (Fig. 1A). When intrinsic apoptosis is initiated, pro-apoptotic BH3-only proteins (e.g. PUMA, NOXA) bind to the BCL2 family proteins to release the BAX/BAK1 monomers. The monomeric effectors oligomerize to form pores in the MOM and trigger an apoptotic cascade (27). MOM permeabilization is often considered to be the “point-of-no-return” for cell death.

**Figure 1.**
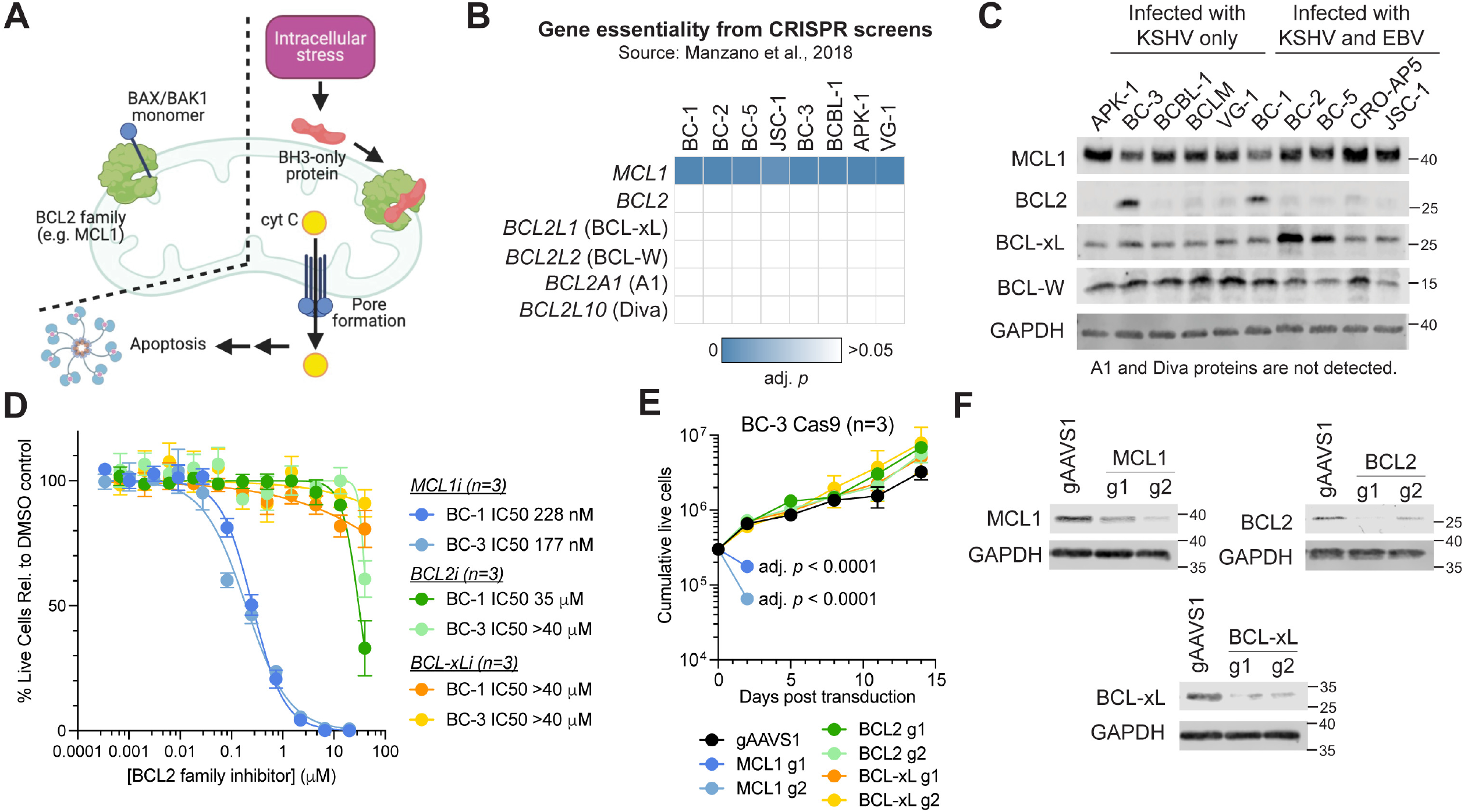
***A***. The BCL2 family proteins (including MCL1) block intrinsic apoptosis by sequestering activated BAX or BAK1 monomers on the outer membrane of the mitochondria. During intrinsic apoptosis, pro-apoptotic BH3-only proteins inactivate the BCL2 family proteins and release the BAX/BAK1 monomers. Free monomers assemble to create mitochondrial pores and release cytochrome c to initiate the apoptotic cascade in the cytoplasm. ***B***. Only gRNAs targeting MCL1 among the BCL2 family are significantly depleted in all PEL cell lines that were previously screened (26). ***C***. Western blots of the different BCL2 family proteins in a panel of PEL cell lines that are infected with KSHV only or co-infected by KSHV and EBV. A1 and Diva proteins are not detected. ***D.*** Dose-response curves of BC-1 and BC-3 cells treated with MCL1i, BCL2i or BCL-xLi. ***E***. Cumulative live cell plot of BC-3 Cas9 cells that were transduced with lentiviruses expressing gRNAs against AAVS1, MCL1, BCL2, or BCL-xL. MCL1 g1 and g2 cells were not counted beyond day 2 because of low number of live cells. Statistical differences were calculated using two-way ANOVA with Dunnett’s multiple comparison test (n=3). ***F***. Representative Western blots of MCL1, BCL2, BCL-xL, or GAPDH at day 2 from panel E. Samples targeting MCL1 and BCL-xL were ran on the same gel flanking the same gAAVS1 control.

Although the BCL2 family members share the same function in BAX/BAK1 sequestration, PEL cell lines appear to be only addicted to MCL1 (Fig. 1B). In recent years, new roles beyond apoptosis have been described for MCL1. MCL1 has been implicated in mitochondrial fusion, respiration, fatty acid oxidation, and autophagy (28–30). In this study, we investigate the basis for why KSHV-transformed PEL cell lines exhibit selective dependence on only MCL1 among the BCL2 gene family. Here, we show that the main essential function of MCL1 in PEL cells is to block intrinsic apoptosis and that non-canonical roles are unlikely. PEL cell lines likely depend on MCL1 because it is expressed at 15-60-fold higher levels than the other BCL2 family members. Altering the expression ratios by over-expression of the other BCL2-related proteins, including the KSHV BCL2 homolog, rescues cells from intrinsic apoptosis induced by an inhibitor of MCL1. This suggests that endogenous expression levels of the BCL2 proteins determine the selective addiction to MCL1 in KSHV-transformed PEL cell lines.

## RESULTS

### Primary effusion lymphoma cells are selectively addicted to MCL1

We previously identified host gene addictions of KSHV-transformed PEL cell lines using genome-wide CRISPR screens (26). Our work showed that these tumor cells require high expression of the oncogene MCL1 for survival. Although the BCL2 family proteins share similar roles in preventing intrinsic apoptosis (Fig. 1A), none of the other BCL2 family genes scored in our CRISPR screens (Fig. 1B). To exclude the possibility that the selective addiction to MCL1 is due to the absence of the other BCL2 family proteins, we performed Western blotting for endogenously expressed BCL2, BCL-xL, BCL-W, A1, Diva, or MCL1 proteins in a panel of PEL cell lines (Fig. 1C). This panel included five cell lines that are co-infected with EBV (BC-1, BC-2, BC-5, CRO-AP5, and JSC-1) and five that are only infected with KSHV (APK-1, BC-3, BCBL-1, BCLM, VG-1). Our immunoblots show that PEL cell lines uniformly express BCL-xL, BCL-W, and MCL1. In contrast, BCL2 expression is detectable only in BC-1 and BC-3 cells, while we do not detect A1 or Diva in any of the cell lines. Similar expression patterns of MCL1, BCL2, and BCL-xL were recently reported in additional PEL cell lines (31).

To confirm the selective dependence of PEL cell lines to MCL1, we treated BC-1 or BC-3 cells with small molecule inhibitors that are specific for MCL1 (S63845, “MCL1i”), BCL2 (ABT-199/venetoclax, “BCL2i”) or BCL-xL (WEHI-539, “BCL-xLi”). These are not expected to cross-inhibit the other BCL2 members. No BCL-W-specific inhibitors have been developed to date. Consistent with our previous work (26), BC-1 and BC-3 cells are sensitive to MCL1i with an estimated half maximal inhibitory concentration (IC50) of 177-228 nM (Fig. 1D). In contrast, both cell lines are resistant to BCL2i and BCL-xLi. These results are consistent with a study showing that PEL cell lines are sensitive to another MCL1i inhibitor AZD-5991 but are resistant to the dual BCL2/BCL-xL inhibitor ABT-263 (31).

To further demonstrate that PEL cell lines selectively depend on MCL1 but not the other BCL2 family members, we targeted the endogenously expressed MCL1, BCL2, or BCL-xL for pooled CRISPR knockout (KO) in BC-3 Cas9 cell lines. Our efforts to target BCL-W for CRISPR KO were unsuccessful. Similar to what we previously observed (26), pooled MCL1 KO cells significantly decreased cell viability by 70-90% as early as two days after transduction of lentivirus encoding MCL1 guide RNAs (gRNAs) (Figs. 1E, 1F). In contrast, pooled BCL2, BCL-xL, and BCL-W cells had similar viability compared to a negative control which targets the *AAVS1* safe harbor locus (gAAVS1) for two weeks. In sum, our data indicate that PEL cell lines express the other BCL2 family proteins, but are selectively addicted to MCL1.

### BAX/BAK1 double knockout cell lines

Roles beyond inhibiting apoptosis have emerged for MCL1, including facilitating mitochondrial respiration and fatty acid oxidation (28–30). We thus explored whether MCL1 may have critical non-canonical functions in PEL cells that necessitate MCL1 expression, but not the expression of other BCL2 proteins, in addition to its canonical role in inhibiting intrinsic apoptosis.

Loss-of-function studies on the non-canonical roles of MCL1 pose a technical hurdle because these are confounded by effects from cell death. To address this issue, we derived clonal BAX and BAK1 double knockout (DKO) cell lines from BC-1 (KSHV+EBV+) and BC-3 (KSHV+) cells (Figs. 2A, 2B). This inactivates the intrinsic apoptosis pathway since pore formation by BAX or BAK1 is considered the necessary step to irreversibly initiate intrinsic apoptosis (Fig. 1A).

**Figure 2.**
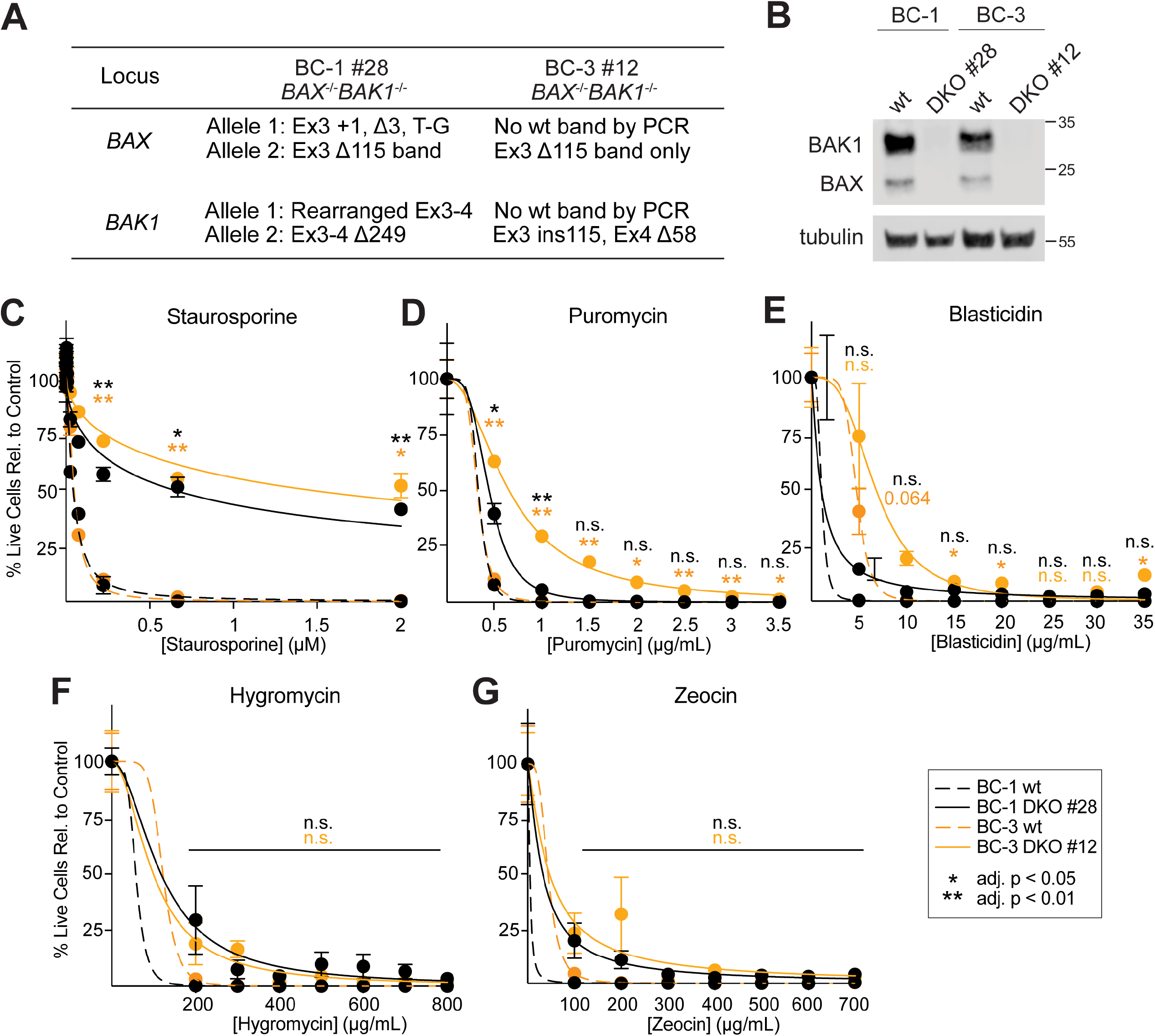
***A.*** Summary of CRISPR edits in BC-1 BAX/BAK1 double knockout (DKO) clone #28 and BC-3 DKO clone #12. ***B.*** Western blots for BAX, BAK1 and tubulin proteins of WT or DKO cells in panel A. . ***C-G.*** Dose response curves of BC-1 or BC-3 cells with BAX/BAK1 wildtype or DKO background upon exposure to different inducers of cell death: *C*. staurosporine, *D.* puromycin, *E*. blasticidin, *F.* hygromycin, or *G*. zeocin. Statistical differences were calculated using two-way ANOVA with Tukey’s multiple comparison post-hoc test (n=3). Adjusted *p* values reflect post-hoc comparisons between the responses of the wt cell line compared to the DKO cell line at the specific time point. Error bars, standard error of mean. n.s., not significant.

We tested whether the DKO mutations successfully inactivate the intrinsic apoptotic pathway after treatment with staurosporine. Staurosporine induces intrinsic apoptosis by broadly inhibiting protein kinases. In parallel, we also tested the sensitivities of the DKO cells to different antibiotics. Dose response curves for these inhibitors show that DKO cell clones are substantially resistant to treatment with staurosporine, but remain sensitive to the antibiotics puromycin, blasticidin, zeocin, and hygromycin (Figs. 2C-G). Our data thus confirm that our DKO cell clones successfully inactivate the intrinsic apoptotic pathway but not other cell death mechanisms.

### Knockout of BAX and BAK1 eliminates the dependency of PEL cells to MCL1

We used the BC-1 and BC-3 DKO cell clones to investigate their dependency on MCL1. If inactivation of MCL1 continues to affect the viability of DKO cells, this would suggest that MCL1 has an essential role in PEL that is independent of intrinsic apoptosis. In contrast, if the DKO cells are unaffected by MCL1 inhibition, then the main essential function of MCL1 is to inhibit BAX/BAK1-mediated cell death.

We therefore treated BC-1 and BC-3 wild type (WT) and DKO cells with increasing doses of the MCL1 inhibitor S63845 (MCL1i) (32). Consistent with our previous work (26) and Fig. 1D, WT BC-1 and BC-3 cells are sensitive to MCL1i (Fig. 3A). However, both DKO cell clones were resistant to MCL1i and viable even at the maximum dose used (20 μM).

**Figure 3.**
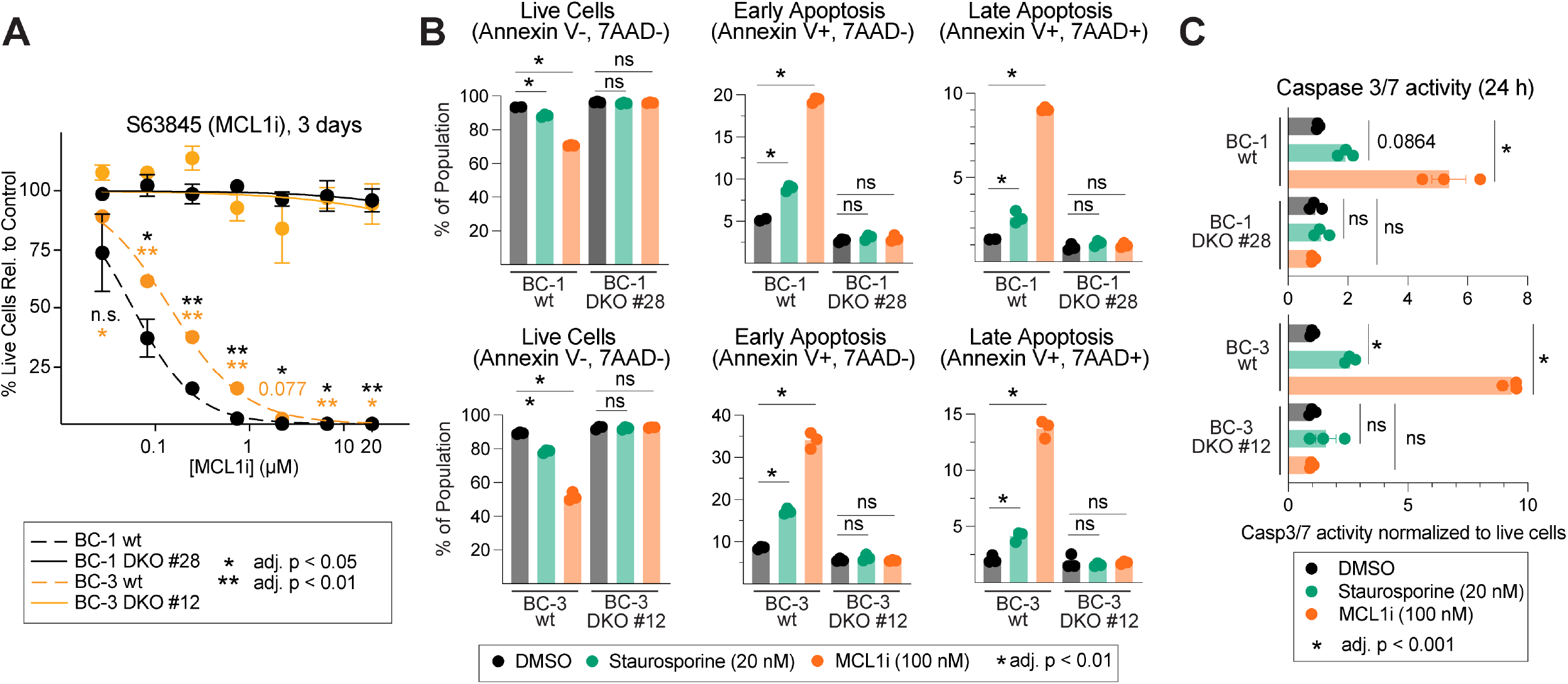
***A***. MCL1i dose response curves of BC-1 or BC-3 cells with BAX/BAK1 WT or DKO background. ***B.*** Distribution of live, early apoptotic or late apoptotic cell populations of BC-1 and BC-3 WT or DKO cells after an overnight treatment of 100 nM MCL1i or 20 nM staurosporine as determined by Annexin V/7AAD staining. ***C***. In parallel, caspase 3/7 activities were measured using a luciferase assay from cells in panel B. Statistical differences were calculated using one-way ANOVA with Tukey’s multiple comparison test (n=3). Error bars, standard error of mean. n.s., not significant.

To confirm whether MCL1i induces apoptosis in PEL cells, we treated the WT and DKO cells with a sublethal dose of MCL1i (100 nM) overnight and measured apoptotic markers using Annexin V/7-AAD staining and Caspase 3/7 Luciferase Assay. We also included a sublethal dose of staurosporine (20 nM) treatment as a positive control. Exposure to MCL1i or staurosporine of BC-1 or BC-3 WT cells decreased live cells between 5-40% compared to the DMSO control (Fig. 3B). This is accompanied by a proportional increase in cells undergoing early and late apoptosis (Fig. 3B) as well as an increase in caspase 3/7 activities (Fig. 3C). In contrast, all of these markers of programmed cell death are completely rescued in BC-1 and BC-3 DKO cells. Thus, apoptotic cell death induced by the pharmacological inhibition of MCL1 is prevented by genetically ablating BAX and BAK1.

Since MCL1i was specifically designed to bind to the BH3-binding groove of MCL1 (32), it is possible that MCL1i induces cell death by only inhibiting the anti-apoptotic role of MCL1 without affecting putative non-canonical functions. We therefore targeted MCL1 for RNAi knockdown in both BC-3 WT and DKO cells using a doxycycline (DOX)-inducible lentiviral shRNA. Total MCL1 protein was reduced by ~70% by quantitative Western blot in both WT and DKO cells using the sh2MCL1 construct after Dox addition (Fig. 4A). This is not seen in the absence of DOX nor in a negative control shRNA construct targeting the *Renilla* luciferase (33). Similar to our previous work (26), MCL1 knockdown lead to a significant decrease in the total number of live cells in BC-3 WT cells as early as two days after DOX treatment (Fig. 4B, dotted red line). This is accompanied by a ~7-fold increase in caspase 3/7 activity (Fig. 4C). In contrast, MCL1 knockdown in BC-3 DKO cells did not substantially affect viability even after 12 days of DOX treatment (Fig. 4B, dotted purple line) nor induce caspase 3/7 activity (Fig. 4C). We note the efficiency of MCL1 knockdown is sustained at this time point as evidenced by reduced MCL1 protein (Fig. 4A, bottom). Taken together, our results from MCL1i treatment or shRNA knockdown of the DKO cells suggest that the essential function of MCL1 in PEL cell lines is to prevent intrinsic apoptosis.

**Figure 4.**
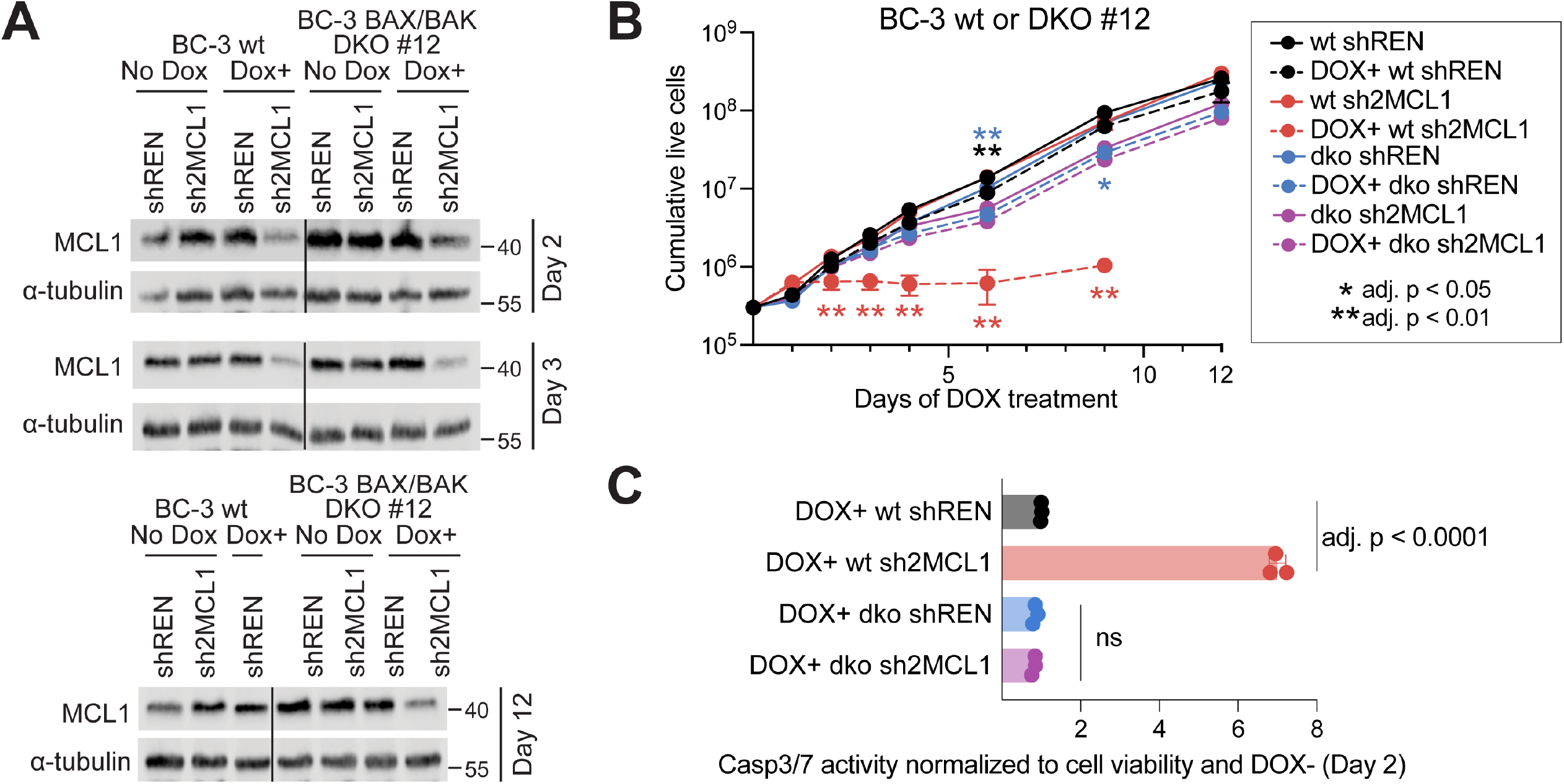
***A.*** Western blots for total MCL1 in BC-3 BAX/BAK wt or DKO #12 cells transduced with lentiviruses expressing a doxycycline (DOX)-inducible shRNA against Renilla luciferase or MCL1. Shown are samples taken after 2, 3, or 12 days of DOX-treatment (1 μg/mL). ***B.*** Cumulative live cell count of BC-3 cells upon DOX treatment (dotted lines). Cell counts were terminated for DOX-treated BC-3 wt cells with sh2MCL1 (dotted red line) at day 9 of DOX-treatment due to significant amount of cell death. Statistical differences were calculated using one- or two-way ANOVA with Tukey’s multiple comparison test (n=3). One-way ANOVA for each time point was instead done for *C*. since two-way ANOVA requires matched data for all time points. Adjusted *p* values reflect post-hoc comparisons between the responses of the wt cell line compared to the DKO cell line at the specific time point or Dox-treated cell line compared to matched untreated cell line (n=3). C. Caspase 3/7 activities were measured using a luciferase assay from cells after 2 days of DOX treatment. Statistical differences were calculated using one-way ANOVA with Tukey’s multiple comparison test (n=3). Error bars, standard error of mean. n.s., not significant.

### Over-expression of BCL2 proteins in PEL cell lines

Our results thus far indicate that the main role of MCL1 in PEL cell lines is to inhibit cell death. However, this does not address why we observe a selective addiction to MCL1 in our CRISPR screens despite co-expression of functionally similar BCL2 proteins. Because a minimum number of pro-survival BCL2 family protein molecules is required to successfully neutralize BAX/BAK1 monomers and prevent pore formation, we hypothesized that the expression ratios of the BCL2 family directly influence the addiction to MCL1. Although we detected BCL-xL and BCL-W proteins in all PEL cell lines and BCL2 in BC-1 and BC-3 (Fig. 1C), Western blotting does not allow for relative quantification between different proteins even in the same sample. We therefore reanalyzed publicly available mRNA-seq datasets from different PEL cell lines including other B cell malignancies to determine the abundance of the BCL2 family transcripts (34). As seen in Figure 5A, endogenous MCL1 is the most highly expressed BCL2 family gene in all six PEL cell lines. In contrast, BCL2, BCL-xL, and BCL-W mRNA levels are only modestly expressed (15- to 60-fold less compared to MCL1) while A1 and Diva mRNAs are not detected. This expression profile reflects the degree of depletion of gRNAs from the CRISPR screens (Fig. 1B) consistent with the model that expression ratios of the BCL2 family may influence the selective addiction to MCL1. Notably, these ratios and magnitude of MCL1 expression in PEL cell lines are similar to what is seen in multiple myeloma cell lines which are also considered to be plasma cell-like. This is consistent with previous reports demonstrating that the PEL and multiple myeloma cell lines have similar gene expression profiles (4–6). These expression patterns of the BCL2 family genes are distinct from those observed in other B cell malignancies that represent different stages of B cell maturation (Fig. 5A). It is possible that the stoichiometry of the BCL2 family members is a function of the stage of B cell differentiation.

**Figure 5.**
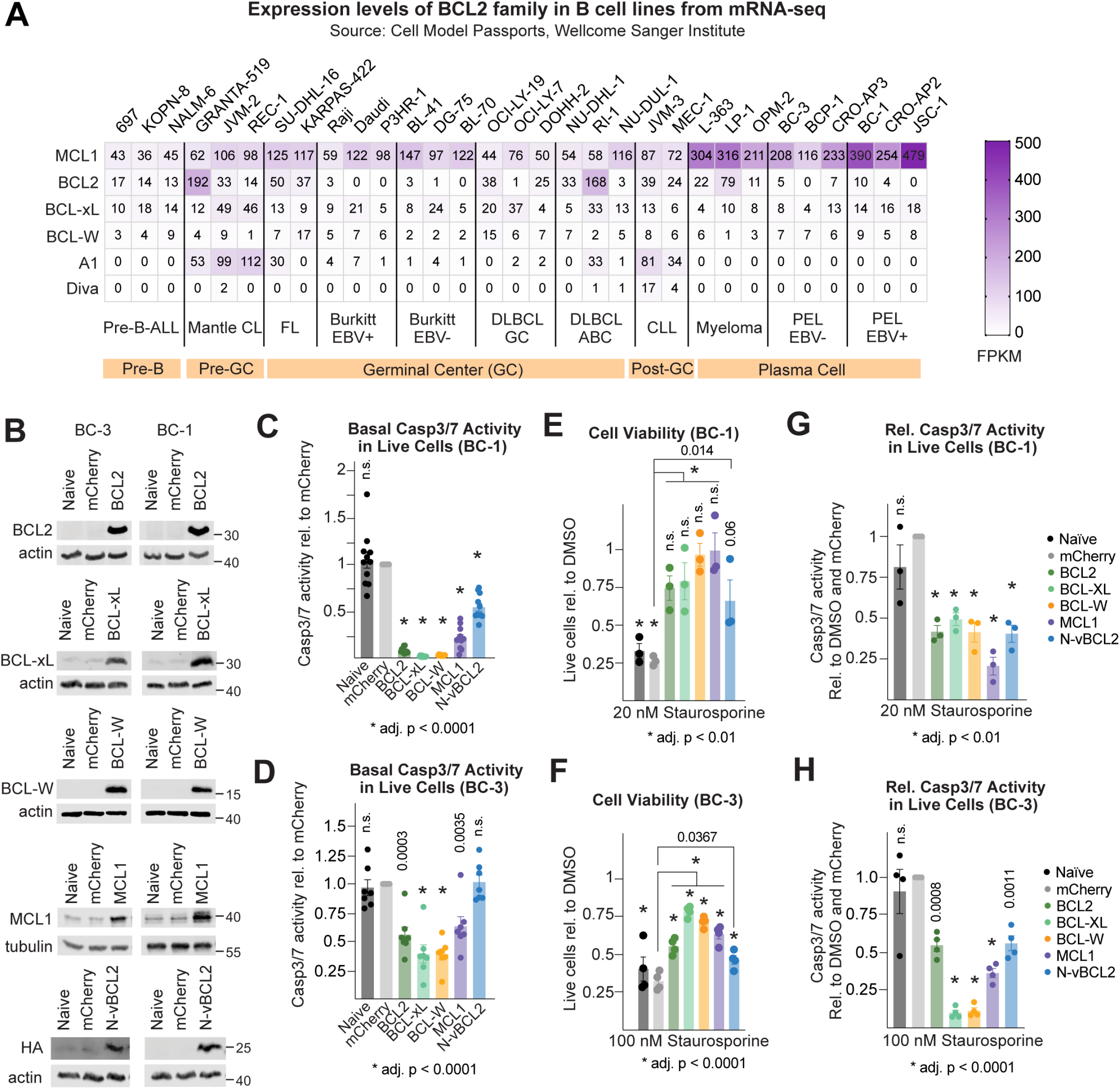
***A***. Normalized expression values (fragments per kilobase of exon per million mapped fragments, FPKM) of BCL2 family transcripts in B cell lines representing different malignancies and stage of B cell differentiation from the Cell Model Passports database (34). These cancers include: pre-B cell acute lymphoblastic leukemia (pre-B-ALL), mantle cell lymphoma (mantle CL), follicular lymphoma (FL), Burkitt lymphoma (EBV-infected or uninfected), diffuse large B cell lymphoma germinal center subtype (DLBCL GC) or activated B cell subtype (DLBCL ABC), chronic lymphocytic leukemia (CLL), multiple myeloma and PEL. ***B.*** Western blots of the BCL2 family members and HA-tagged vBCL2 in BC-1 or BC-3 cells transduced with lentiviruses over-expressing these cDNAs. ***C-D.*** Basal caspase 3/7 activity of BC-1 (n=11) or BC-3 cells (n=7) normalized to total live cells measured by a caspase 3/7 luciferase assay. ***E-F.*** Cell viability measured by CellTiterGlo 2.0 assay or ***G-H.*** relative caspase 3/7 activity in live BC-1 (n=3) or BC-3 cells (n=4) after overnight treatment of 20 nM or 100 nM staurosporine. Statistical differences were calculated using one- or two-way ANOVA with Tukey’s multiple comparison post-hoc test. Adjusted *p* values reflect post-hoc comparisons between the cell lines in response to staurosporine or DMSO unless indicated otherwise. Error bars, standard error of mean. n.s., not significant.

If the skewed expression of MCL1 explains the dependency of PEL cell lines to this BCL2 family member, we expect that changing the ratios of the BCL2 family genes would shift the dependency to the most abundant BCL2 family protein and away from MCL1. We thus transduced BC-1 or BC-3 cells with lentiviruses constitutively expressing the different BCL2 genes under the major immediate-early promoter and enhancer of the human cytomegalovirus at a multiplicity of infection of 5. We chose to only over-express the proteins that are expressed endogenously at the RNA and protein levels: BCL2, BCL-xL, BCL-W, or MCL1. In addition, we also expressed the KSHV homolog of BCL2, vBCL2 (35, 36), that has been dually tagged with FLAG-hemagglutinin (HA) epitopes at the N-terminus (N-vBCL2). Although vBCL2 is a lytic gene and is not expressed in latently infected PEL cells, we were curious if this can functionally substitute for MCL1 in PEL. Western blots confirm that each protein is expressed above endogenous levels in both BC-1 and BC-3 backgrounds (Fig. 5B).

To confirm that the over-expressed proteins are functional in inhibiting intrinsic apoptosis, we induced cell death in these cell lines using staurosporine and measured cellular viability using CellTiterGlo 2.0 and apoptosis using Caspase 3/7 Luciferase Assay. All of the ectopically expressed BCL2 family proteins and vBCL2 reduced basal caspase 3/7 activity in the absence of staurosporine in both BC-1 and BC-3 cells (Figs. 5C, 5D). Reduction of caspase 3/7 activity was consistently stronger in the BC-1 cell lines compared to the BC-3 cell lines. In both cases, over-expression of BCL2, BCL-xL, or BCL-W inhibited basal caspase 3/7 activity the most while MCL1 or N-vBCL2 had modest to no effects. It is possible that this minor effect of MCL1 or N-vBCL2 in decreasing basal caspase 3/7 activity may be due to insufficient over-expression of these proteins.

In the presence of staurosporine, naïve and mCherry-expressing cells had significantly reduced cellular viability after overnight treatment as expected (Figs. 5E, 5F). Cell death was rescued in all instances by all the BCL2 family proteins including N-vBCL2 although to a lesser extent compared to the mCherry control. These effects in cell viability are conversely accompanied by a reduction in caspase 3/7 activities (Figs. 5G, 5H). These experiments demonstrate that the ectopically expressed BCL2 family proteins in BC-1 or BC-3 cells are functional in inhibiting intrinsic apoptosis.

### Over-expression of BCL2 family proteins desensitizes PEL cells to MCL1i-induced apoptosis

We next tested whether over-expression of the BCL2-related proteins in BC-1 or BC-3 can shift the dependency to a different BCL2 family protein and cause to cells to be less reliant to MCL1 for survival. We therefore inactivated MCL1 with MCL1i. Similar to staurosporine, overnight treatment with MCL1i reduced the viability of naïve or mCherry-expressing cells (Fig. 6A). In contrast, ectopic BCL2, BCL-xL, or BCL-W fully protected BC-1 and BC-3 cells against MCL1i toxicity. N-vBCL2- and surprisingly, MCL1-expressing cells were sensitive to MCL1i with only a marginal rescue of cell viability (20-30%) compared to mCherry. These effects in cell viability are likely due to apoptosis as seen with the opposite pattern of caspase 3/7 activity in these cell lines (Fig. 6B). Relative to naïve and mCherry-expressing cells, ectopic BCL2, BCL-xL, or BCL-W inhibited caspase 3/7 activity induced by MCL1i to the same degree in both BC-1 and BC-3. Ectopic N-vBCL2 moderately reduced caspase 3/7 activity while MCL1 had no measurable effect.

**Figure 6.**
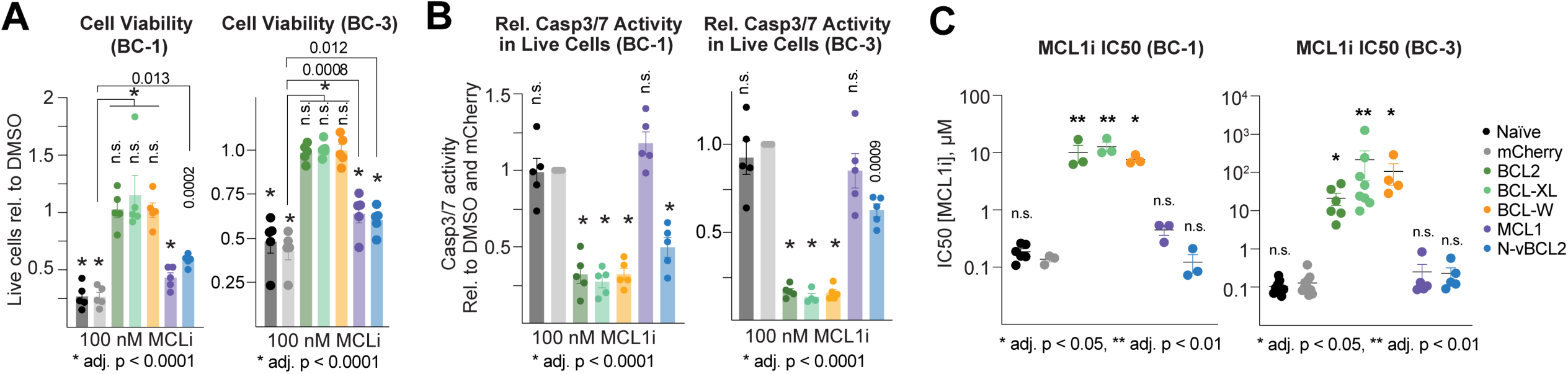
***A.*** Cell viability measured by CellTiterGlo 2.0 assay or ***B.*** relative caspase 3/7 activity in live BC-1 or BC-3 cells over-expressing BCL2 family cDNAs after overnight treatment of 100 nM MCL1i. Statistical differences were calculated using one- or two-way ANOVA with Tukey’s multiple comparison test (n=5). Adjusted *p* values reflect post-hoc comparisons between the cell lines in response to staurosporine or DMSO unless indicated otherwise. ***C.*** IC50 values of MCL1i in BC-1 or BC-3 cells. For BC-1, one-way ANOVA with Tukey’s multiple comparison test was used for statistical tests (n≥3). Since BC-3 cell lines over-expressing BCL2, BCL-xL or BCL-W were highly resistant to MCL1i even at high doses, IC50 values derived from these incomplete sigmoidal dose-response curves had low confidence intervals. Thus, non-parametric Kruskal-Wallis post-test was used. Adjusted *p* values reflect post-hoc comparisons between the cell lines to the mCherry control. Error bars, standard error of mean. n.s., not significant.

Finally, we treated the cells with varying doses of MCL1i to calculate their relative sensitivities to this inhibitor. Consistent with what we observed in Figs. 6A and 6B, over-expression of BCL2, BCL-xL or BCL-W desensitized cells from MCL1i and increased the IC50 values by 100- to 1000-fold (Fig. 6C). The N-vBCL2 construct did not alter the over-all sensitivities of BC-1 or BC-3 cells to MCL1i. While ectopic MCL1 increased the average MCL1i IC50 by ~2- (BC-3) to 3-fold (BC-1), these effects were not statistically significant. In sum, our data demonstrates that expression levels of the BCL2 family genes is the major determinant for the dependency to MCL1 by PEL cell lines.

## DISCUSSION

In this report, we explore why KSHV-transformed PEL cell lines exhibit selective addiction to the oncogene MCL1, despite having shared functions with the related BCL2 family proteins. While new roles beyond those shared with the other BCL2 family members have been attributed to MCL1, we show that the essential function of MCL1 in PEL cell lines is to prevent intrinsic apoptosis. This selective addiction to MCL1 is explained by the 15- to 60-fold stoichiometric excess of MCL1 expression relative to other BCL2-related proteins as demonstrated by mRNA-seq and functional rescue from MCL1i-induced cell death by over-expression of any of the BCL2 family proteins.

The failure to rescue MCL1i-induced apoptosis by the MCL1 or N-vBCL2 constructs may be due to the degree of ectopic expression achieved in these cells. It is possible that the total amount of BCL2 family proteins in any of these cases did not reach the minimum required to neutralize all available MCL1i molecules.

It is not clear why BCL2 family expression is skewed towards MCL1 in PEL cell lines. We speculate that this may be a consequence of the plasma cell-like B cell differentiation stage of PEL tumors (4–6). Similarly, Fig. 5A and the Cancer Dependency Map (37) shows that multiple myeloma cell lines, which are considered to be of plasma cell lineage, have high MCL1 expression and are selectively dependent on MCL1. In contrast, cell lines derived from B cell malignancies representing earlier B cell stages have distinct expression patterns where the other BCL2 family members are expressed at higher levels (Fig. 5A). Studies done in mice by the Tarlinton group indicate that the requirements for the Bcl2 family members during B cell development are dynamic and complex. It was shown that Mcl1 but not Bcl-xL is required for the formation but not persistence of germinal center B cells and memory B cells using conditional knockout models (38). This points to a transient dependence of antigen-specific B cells on Mcl1 for competition within germinal centers during maturation but not for their survival after their establishment. Mcl1, however, is indispensable for the survival of plasma cells in culture and in vivo in a tamoxifen-inducible Mcl1 knockout mouse model in a cell intrinsic manner (39). In contrast, inducible knockout of Bcl-xL or treatment with ABT-737, a preferential inhibitor of BCL2 with modest effects on Bcl-xL and Bcl-W, have no effect in the survival of existing plasma cells in culture or in vivo, respectively (39, 40). They propose that plasma cells require high levels of Mcl1 along with other factors to provide an intrinsic survival potential in combination with extrinsic survival signals from tissue-specific niches (e.g. secondary lymphoid tissues or bone marrow) to ensure long-lived plasma cells (41). The requirement for the high expression of MCL1 in plasma cells may be driven by master transcription factors such as IRF4 that occupy lineage-specific super-enhancer regulatory regions. Indeed, we and others have identified a super-enhancer element around the MCL1 locus bound by IRF4 in PEL cell lines (21, 42, 43).

Upon activation and during maturation, B cells undergo dramatic changes to support their effector functions in immune defense (Reviewed in 44). A large part of the plasma cell differentiation program is the remodeling and expansion of the endoplasmic reticulum (ER) to prepare for immunoglobulin production and secretion (45). Nutrient transporters and metabolic enzymes are upregulated to sustain antibody production and organelle maintenance by the lineage transcription factor Blimp-1 (46). High expression of MCL1 may thus act as a buffer from this increased ER stress and metabolic load to ensure the survival of these B cells. Knockdown of the glucose transporter GLUT4 in multiple myeloma cell lines induced apoptosis by decreasing MCL1 mRNA expression (47), thereby supporting this link between metabolism and apoptosis.

In the case of PEL, the infected tumor cells neither express nor secrete immunoglobulin molecules. However, recent work from the Gottwein laboratory shows that PEL cell lines need to constitutively counteract death signals originating from the ER and Golgi compartments in the absence of any exogenous trigger (48). Consistent with this, PEL cell lines have elevated expression of genes involved in the unfolded protein response, often triggered by ER stress (4). These suggest that the ER and Golgi bodies in PEL cells are also under stress despite the absence of antibody production. In line with this, KSHV is known to activate ER stress responses during lytic reactivation (49, 50). Therefore, KSHV needs to counter these ER, Golgi and metabolic stresses by possibly reinforcing or amplifying existing survival pathways in plasma cells such as MCL1. More recently, the latency-associated nuclear antigen (LANA) was shown to indirectly enhance the steady state levels of the MCL1 protein by binding to the E3 ubiquitin ligase FBW7 and preventing MCL1 degradation (51). As mentioned above, although we find binding sites for the lineage transcription factor IRF4 on the MCL1 super-enhancer in PEL cells, we recently reported that the B cell-specific latency gene vIRF3 largely co-occupies similar sites as IRF4, including the MCL1 super-enhancer and promoter (21). This suggests that vIRF3 may function to enhance MCL1 expression. We are currently investigating how vIRF3 regulates MCL1 expression at the transcriptional level.

A parallel relationship between lineage-specific expression and dependence to the BCL2 family proteins is also seen in lymphatic endothelial cells (LECs) which are thought to be candidates for the cellular origin of Kaposi’s sarcoma spindle cells (52). In this case, LECs express and depend on high levels of BCL-xL instead of MCL1 for survival. This dependency dramatically increases during KSHV infection (Dr. Michael Lagunoff, personal communication) which supports the model where the virus reinforces pro-survival pathways already present in the cell.

The selective dependence of PEL cell lines to MCL1 makes it an attractive target for therapy. We have previously shown that the MCL1 protein is highly expressed in primary tumors from patients (26). Candidate MCL1-selective drugs are now in various stages of clinical trials to treat hematologic malignancies (53) and have the potential to be repurposed to treat PEL. Venetoclax, the first approved BH3 mimetic drug, has over-all response rates of ~80% in relapsed and/or refractory chronic lymphocytic leukemia (54) and ~75% in relapsed and/or refractory mantle cell lymphoma (55) with a durable response even after ~3 years (56). However, other cancers like acute myelogenous leukemia exhibit a lower response rate to venetoclax (19%) where non-responsive patients had an increase in BCL-xL or MCL1 expression (57). Our data similarly shows that over-expression of any of the BCL2 family proteins or loss of BAX or BAK1 leads to resistance to MCL1 inhibitors. Thus, care must be taken when testing MCL1 inhibitors in PEL patients to avoid resistance. One strategy is using high but tolerable doses of MCL1 drugs to minimize selection for resistant tumor cells. Another approach is to combine this with other regimens that do not act on the same pathway. These strategies need to be explored to provide the most effective treatment for KSHV-induced PEL.

## MATERIALS AND METHODS

### Cells

Cells were cultured as described (21). HEK293T cells were cultured in Dulbecco’s modified Eagle medium with L-glutamine and 4.5 g/L glucose (Corning Life Sciences, Tewksbury, MA) supplemented with 10% Serum Plus II medium (Millipore-Sigma, Burlington, MA) and 10 μg/ml gentamicin (Gibco, Waltham, MA). BC-1 and BC-3 cell lines were grown in RPMI 1640 (Corning) supplemented with 10 μg/ml gentamicin (Life Technologies, Carlsbad, CA), 0.05 mM β-mercaptoethanol (Sigma-Aldrich, St. Louis, MO), and 10% (BC-1) or 20% (BC-3) Serum Plus II medium. All PEL cell lines were grown at 37°C with 5% CO_2_ and maintained at log phase between 2×10^5^ and 1×10^6^ cells/mL.

### Primers and synthetic genes

All primers were purchased from Integrated DNA Technologies (Coralville, IO). Synthetic genes for eGFP fragment, shRNA cassettes and MCL1 cDNA were synthesized by Twist Bioscience (South San Francisco, CA). Sequences and primer combinations for cloning are listed in Tables 1 and 2.

**Table 1.**
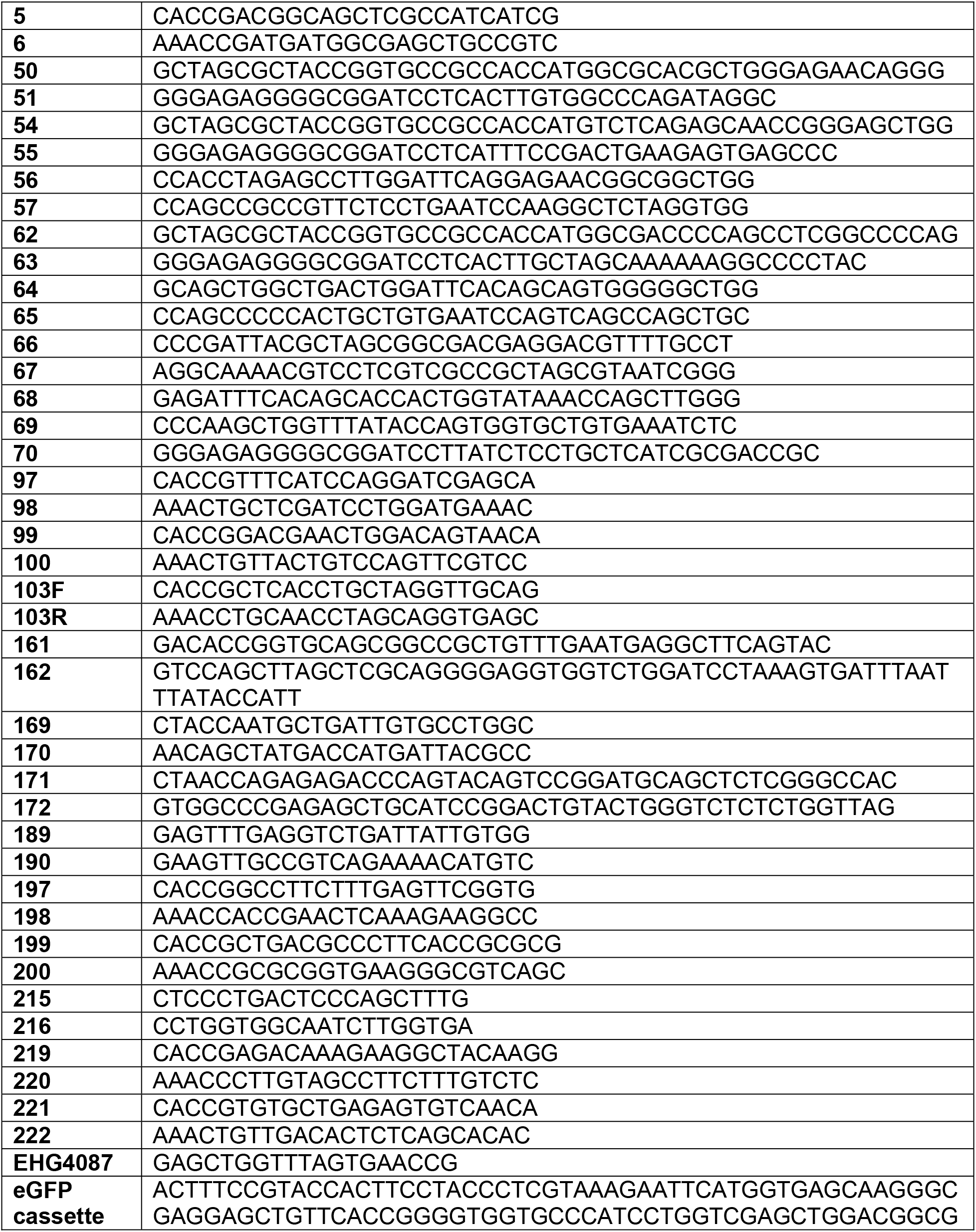
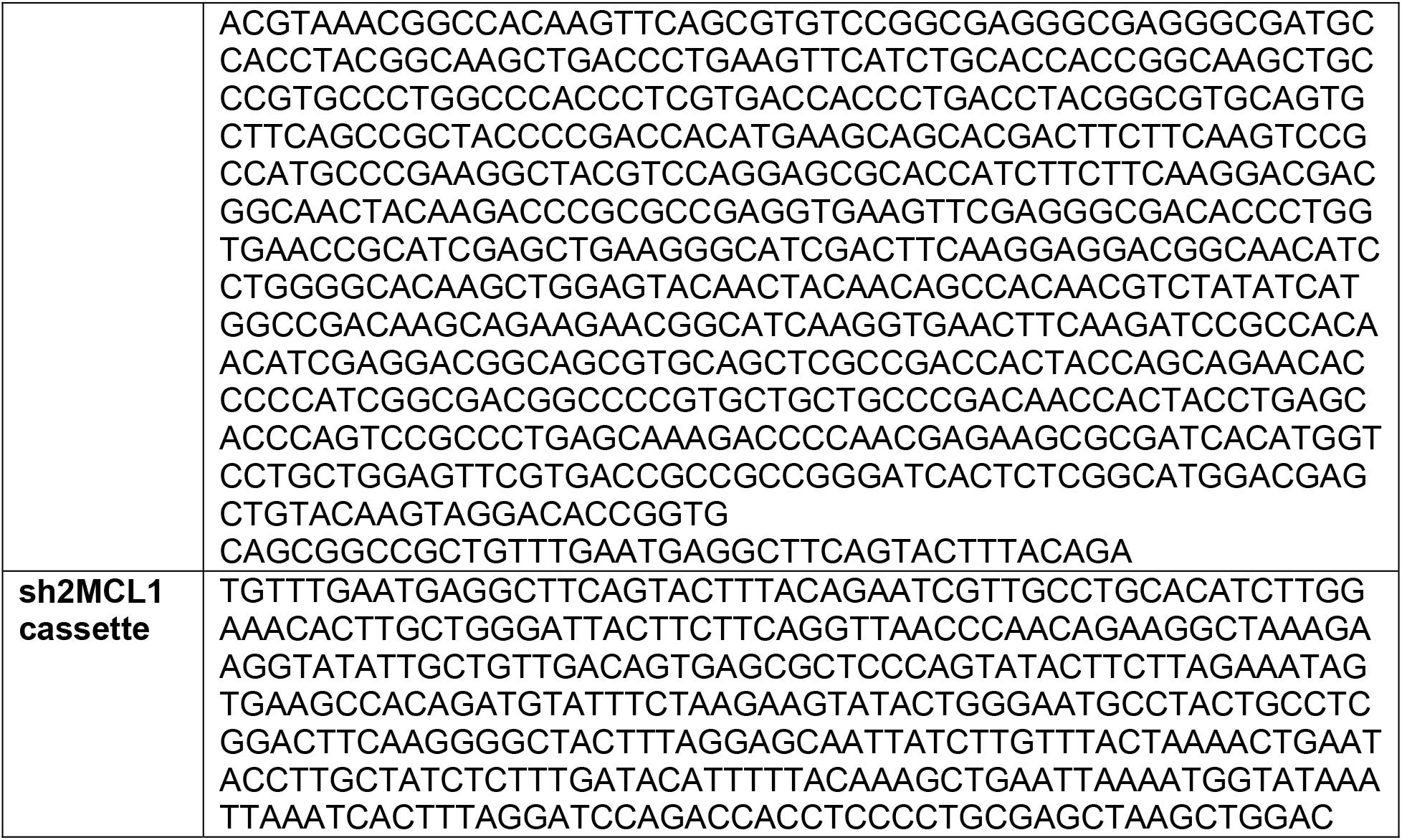
Primer sequences.

**Table 2.** Combinations of primers used for cloning constructs.

| Construct | PCR | Primers | Template | Source |
| --- | --- | --- | --- | --- |
| pLV-SIN-TetOne-ZeoR (pSTZ) | 1 | 169, 171 | pLVX-TetOne-Zeo | (58) |
|  | 2 | 170, 172 |  |  |
|  | Overlap | 169, 170 | PCR 1 and 2 |  |
| shREN | N/A | 161, 162, | pZIP-ZsGreen-T2A-Hyg-shRen.713 | (48) |
| BCL2 | N/A | 50, 51 | pCDNA3 HA-BCL2 | (59) |
| BCL-xL | 1 | 54, 57 | pcDNA3 HA-BCL-xL |  |
|  | 2 | 55, 56 |  |  |
|  | Overlap | 54, 55 | PCR 1 and 2 |  |
| BCL-W | 1 | 62, 65 | pLX304 BCL2L2 | DNASU |
|  | 2 | 63, 64 |  |  |
|  | Overlap | 62, 63 | PCR 1 and 2 |  |
| vBCL2 N-FLAG-HA | 1 | EHG4087, 67 | pLC-FLAG-HA-vIRF3-P2A-hyg | Manzano & Gottwein, unpublished |
|  | 2 | 66, 69 | BC-1 genomic DNA |  |
|  | 3 | 68, 70 | BC-1 genomic DNA |  |
|  | 1+2 | EHG4087, 69 | PCR 1 and 2 |  |
|  | Final | EHG4087, 70 | PCR 1+2 and 3 |  |
| BAX g2 | Anneal | 97, 98 |  |  |
| BAX g3 | Anneal | 99, 100 |  |  |
| BAK1 g1 | Anneal | 5, 6 |  |  |
| BAK1 g2 | Anneal | 103F, 103R |  |  |
| BCL2 g1 | Anneal | 197, 198 |  |  |
| BCL2 g2 | Anneal | 199, 200 |  |  |
| BCL-W g1 | Anneal | 219, 220 |  |  |
| BCL-W g2 | Anneal | 221, 222 |  |  |

### shRNAs

A self-inactivating version of the pLVX-TetOne-Zeo lentiviral vector (58) (a gift from Dr. Britt Glaunsinger) was first made via PCR mutagenesis. A deletion in the U3 region of the 3’LTR was introduced using primer pairs 169/171 and 172/170. The PCR products were fused with primers 169/170 and cloned into pLVX-TetOne-Zeo using the KpnI and NheI restriction sites (New England Biolabs, Ipswich, MA). The new vector is renamed pLVX-SIN-TetOne-Zeo (pSTZ) to denote the self-inactivating deletion (SIN).

shRNAs were designed using the SplashRNA algorithm (33) using default parameters (http://splashrna.mskcc.org). sh2MCL1 was synthesized as a gene fragment (Twist Bioscience) while shREN was amplified from pZIP-ZsGreen-T2A-Hyg-shRen.713 (48). These shRNA cassettes were assembled in the 3’UTR of a synthetic eGFP gene (Twist Bioscience) via two-fragment Gibson assembly into the pSTZ backbone linearized by EcoRI and BamHI (NEB).

### BCL2 cDNAs

cDNAs for human BCL2, BCL-xL, and BCL-W were amplified from pCDNA3-HA-BCL2, pCDNA-HA-BCL-xL (59) (gifts from Dr. Timothy Chambers), and pLX304-BCL2L2 (Clone # HsCD00440394 from Human ORFeome Collection from DNASU (60)). vBCL2 sequence was amplified from BC-1 genomic DNA. The Flag-HA dual tag cassette was amplified from pLC-FLAG-HA-vIRF3-P2A-hyg (Manzano and Gottwein, unpublished). Internal restriction sites were repaired using PCR mutagenesis. Details on primer sequences and primer combinations for each PCR step are listed in Tables 1 and 2.

### Lentiviruses

The day prior to transfections, 4-5×10^6^ 293T cells were plated in 10 cm dishes. Lentiviral transfer vectors (5 μg) were co-transfected with pMD2.G (2.44 μg) and psPAX2 (3.74 μg) packaging vectors and 39.13 μL 1 mg/mL polyethylenimine (Sigma Catalog #408727) in 500 μL OptiMEM (Gibco). After 4-6 hours, media was replaced with complete RPMI medium supplemented with 10% Serum Plus II medium. Supernatants containing lentiviruses were harvested after three days by filtering through a 0.45 μm pore membrane.

Lentiviral titers were determined by functional titration. Increasing volumes of supernatant containing lentiviruses were added to 1×10^5^ naïve BC-1 or BC-3 cells in 0.5 mL media with 0.5 μL 8 μg/mL polybrene. The day after, appropriate antibiotics were added to select for infected cells. Number of surviving cells was compared to uninfected cells to determine infection rates and viral titers. Final lentiviral transductions were performed in 4×10^5^ BC-1 or BC-3 cells in 2 mL media with 8 μg/mL polybrene at defined MOIs.

### Generation of BAX/BAK1 DKO cells

Two gRNAs targeting the human *BAX* or *BAK1* gene were chosen from the genome-wide Brunello CRISPR library to increase the efficiency of editing in all alleles (61). Based on these gRNAs, primer pairs were designed to create dsDNA duplexes with compatible BbsI overhangs. These primer pairs were assembled by heat-denaturation and annealing in Buffer 3.1 (NEB), then cloned into BbsI-linearized pX458 (62) (a gift from Dr. Feng Zhang, Addgene #48138).

BC-1 and BC-3 cells were adjusted to 2×10^5^ cells/mL the day before transfection. On the day of transfection, 1×10^6^ BC-1 or BC-3 cells were pelleted in 50 mL tubes at 90g, for 10 min. Each pellet was carefully resuspended in 20 μL of SF Cell Line Nucleofector Solution with Supplement 1 (Lonza Cat. #V4XC-2032, Basel, Switzerland) and 1.5 μL of pX458 plasmids (150 ng each of BAX g2, BAX g3, BAK1 g1, BAK1 g3; 600 ng total). Cells transferred to Nucleocuvette strips and pulsed with program CA-189 using the 4D-Nucleofector X (Lonza). Transfection reactions were allowed to recover at room temperature for 10 min. Afterwards, 80 μL of pre-warmed complete media was added. Cells in the Nucleocuvette strips were incubated at 37°C for 10 min. Finally, cells were collected with an additional 120 μL of pre-warmed complete media and transferred to a round-bottom 96-well plate. Cells were expanded for 5-7 days to at least 2×10^6^ total cells. Two additional rounds of nucleofection were performed as above to ensure efficient editing in all four alleles.

After a week of recovery from the third nucleofection, cells were individually cultured in round bottom 96-well plate using dilution cloning. Clones were screened for successful editing. We first triaged the clones based on the absence of full length BAX and BAK1 proteins by Western blotting. Next, we further screened clones using genotyping primers (189 and 190 for *BAX*; 215 and 216 for *BAK1*). We would expect that simultaneous cutting of the two guides in an allele will lead to ~100-400 bp deletions that can be distinguished from wt or single cut alleles by PCR. While this assay does not discriminate between unedited from single editing events, this allows us to limit our final validation to those with at least one allele deleted. Finally, we confirmed frameshift indels by amplifying the targeted loci for Sanger or Illumina sequencing. Illumina sequencing and analysis were performed by the Massachusetts General Hospital Center for Computational and Integrative Biology DNA Core Facility.

### CRISPR targeting of BCL2 family

Clonal BC-1 Cas9 or BC-3 Cas9 cells (26) were transduced with lentiviruses encoding gRNAs targeting the BCL2 family genes at MOI 1. gRNAs targeting BCL2 or BCL-W were cloned in pLentiGuide (Addgene #117986) that was linearized with BsmBI (NEB). gRNAs for gAAVS1, BCL-xL and MCL1 have been previously cloned and described (26). Live cells were measured every 2-3 days on the BD FACS Celesta Flow Cytometer (BD Biosciences, Franklin Lakes, NJ) using a known amount of spike-in AccuCount Blank particles (Spherotech, Lake Forest, IL, Cat. #ACBP-50-10). Data analysis was performed using FlowJo v10.8.1 software (BD Biosciences). Cells were split to 3 × 10^5^ cells/mL to ensure that all cells were continuously dividing.

### Western blots

Cells were washed with phosphate buffered saline and lysed with RIPA lysis buffer (50 mM Tris, 150 mM NaCl, 1% IGEPAL CA-630, 0.5% sodium deoxycholate, 0.1% sodium dodecyl sulfate, pH 7.4) with protease inhibitor cocktail III (Calbiochem, EMD Millipore). Lysates were incubated on ice for 10 minutes and clarified by spinning at 12,000 g for 10 minutes. Total protein was quantified the Pierce BCA Protein Assay Kit (Thermo Fisher).

Equivalent amounts of total proteins were run in Bolt 4%-12% Bis-Tris gradient gels (Thermo Fisher Scientific) or 15% tris-glycine-SDS acrylamide gels and transferred to 0.2 μm-pore size nitrocellulose membranes in Towbin buffer (25 mM Tris, 192 mM glycine, 20% methanol, pH 8.3) for 90 min at 400 mA. Membranes were blocked with a 1:3 dilution of Intercept (TBS) Blocking Buffer (Li-Cor Biosciences, Lincoln, NE) in TBS for 1 h at room temperature. Membranes were incubated with primary antibodies diluted 1:3000 in blocking buffer–0.1% Tween 20 overnight at 4°C. Primary antibodies were purchased from Cell Signaling Technology (Danvers, MA): anti-BCL2 clone D55G8, anti-BCL-xL clone 54H6, anti-BCL-W clone 31H4, anti-MCL1 clone D2W9E, anti-HA clone C29F4, anti-A1 clone D1A1C, anti-Diva (Catalog # 3869), anti-BAK clone D4E4, and anti-BAX clone D2E11; or Sigma: anti-beta-actin clone AC-15 and anti-alpha-tubulin clone B-5-1-2; or Santa Cruz Biotechnology (Dallas, TX): anti-GAPDH clone 0411.

Immunoblots were washed five times with TBST for 5 min each then incubated for 1 hour with IRDye 800 CW-conjugated secondary antibodies (Li-Cor Biosciences) diluted in TBS (1:20,000). Membranes were finally washed four times with TBST and once in TBS. Western blots were imaged on the Odyssey Fc or M Imager (Li-Cor Biosciences).

### Dose Response Curves

Serial dilutions of the different inhibitors/drugs were added to each well containing 5 × 10^3^ BC-1 or BC-3 cells in a U-bottom 96-well plate. At designated time points (2 days for puromycin (Cayman Chemicals, Ann Arbor, MI); 3 days for MCL1i (MedChemExpress, Monmouth Junction, NJ), ABT-199 (Tocris Bioscience, Minneapolis, MN), WEHI-539 (APExBIO, Houston, TX) and staurosporine (APExBIO); 6 days for blasticidin (Gibco), zeocin (Invivogen, San Diego, CA), and hygromycin (Gibco)), 20 μL aliquots of the cells were used to measure cell viability using 20 μL CellTiter-Glo 2.0 Cell Viability Reagent (Promega). Luminescence was read using Synergy H1 microplate reader (Biotek, Winooski, VT) with an integration time of 2 sec. Raw luminescence readings were normalized as a percentage of the DMSO control. Normalized values were used to calculate the IC50 using the “[inhibitor] vs. normalized response (variable slope)” function in Prism 9 (GraphPad, San Diego, CA).

### Caspase 3/7 Assay

2 × 10^4^ BC-1 or BC-3 cells were seeded in each well of a U-bottom 96-well plate and treated with staurosporine (20 nM for BC-1, 100 nM for BC-3), 100 nM MCL1i, or DMSO. These concentrations were chosen because they approximate the IC50 values of each drug for each cell line. After 24 hours, 20 μL of cells were mixed with 20 μL of the CellTiter-Glo 2.0 Cell Viability Reagent or Caspase-Glo 3/7 Assay substrate (Promega). Luminescence was measured after 2 min incubation for CellTiter-Glo 2.0 or 30 min incubation for Caspase-Glo 3/7 Assay using the Synergy H1 microplate reader as above. Raw luminescence values for caspase 3/7 activities were sequentially normalized to the matched raw CellTiter-Glo 2.0 readings, then to the DMSO controls and finally, to the mCherry-expressing cell line.

### Annexin V/7-AAD staining

BC-1 or BC-3 wt or DKO cells (4 × 10^5^ cells/mL, 3 mL plated in 6-well plate) were treated overnight with DMSO, 20 nM staurosporine, or 100 nM MCL1i. 7 × 10^5^ live cells were pelleted and washed twice with wash buffer (2% bovine serum albumin in PBS). Cells were resuspended in 100 μL Annexin V Binding Buffer (Biolegend, San Diego, CA). Cells were stained for 15 min in the dark with 5 μL APC Annexin V (Biolegend) and 5 μL 7-AAD Viability Staining Solution (Biolegend). After incubation, 400 μL of Annexin V Binding Buffer was added and the cells were analyzed on a flow cytometer. In parallel, cellular viability and caspase 3/7 activities were measured as described above from 20 μL aliquots.

## ACKNOWLEDGMENTS

We thank Logan Moorhead for initial help with engineering the DKO cell lines, Dr. Eva Gottwein for the pZIP-ZsGreen-T2A-Hyg-shRen.713 and pLC-FLAG-HA-vIRF3-P2A-hyg plasmids, Dr. Britt Glaunsinger for the pLVX-TetOneZeo plasmid, and Dr. Timothy Chambers for the BCL-xL and BCL2 expression vectors. We thanks Drs. Eva Gottwein, Youssef Aachoui, Changhoon Oh, and J. Craig Forrest for helpful discussion and feedback on the manuscript.

This work is supported by a Transition Career Development Award K22 CA241355 from the National Cancer Institute of the National Institutes of Health (NIH) to M.M. Additional support was provided by the Center for Microbial Pathogenesis and Host Inflammatory Responses program grant P20 GM103625 from the National Institute of General Medical Sciences (NIGMS) of the NIH, Winthrop P. Rockefeller Cancer Institute at UAMS, and an equipment grant from the UAMS Office of the Vice Chancellor for Research and Innovation. J.G. is a recipient of the Undergraduate Summer Research Fellowship from the Arkansas IDeA Network of Biomedical Research Excellence (INBRE) funded by program grant P20 GM103429 from NIH/NIGMS. The content is solely the responsibility of the authors and does not necessarily represent the official views of the funding bodies. The authors declare no conflicts of interests.

